# Conserved switch genes regulate a novel cannibalistic morph after whole genome duplication

**DOI:** 10.1101/2023.08.22.554244

**Authors:** Sara Wighard, Hanh Witte, Ralf J. Sommer

## Abstract

Developmental plasticity facilitates morphological and behavioural novelty, but associated regulatory mechanisms remain elusive. Nematodes have emerged as a powerful model to study developmental plasticity and its evolution. Here, we show the predatory nematode *Allodiplogaster sudhausi* evolved an additional third mouth morph, concomitant with whole genome duplication (WGD) and a strong increase in body size. The three mouth morphs are induced by different diets; bacteria, fungi and nematodes. CRISPR experiments indicate that regulation of the third morph involves co-option of a conserved developmental switch gene, which through WGD resulted in two mouth-form regulators. Gene dosage studies revealed a diverged role of these developmental switches, with functional redundancy and quantitative effects in the two mouth-form decisions, respectively. The third morph is cannibalistic and kills kin, whereas the other two morphs do not. Thus, the recent evolution of a new morph relies on pre-existing regulatory mechanisms and adds behavioural and social complexity.

**One-Sentence Summary:** Experimental genetics in a nematode reveals a key role for developmental plasticity in the evolution of nutritional diversity

## Main Text

Novel morphological structures and behaviors are often observed in developmentally plastic systems, including ant worker castes and predatory spadefoot toads (*1, 2*). However, the regulatory processes and molecular mechanisms that lead to novel traits remain elusive (*3*). Does the evolution of novelty require large-scale genomic alterations? Is it controlled by pre-existing or novel regulatory genes and mechanisms? And what type of environmental variations, *e.g*. changes in diet, are involved in such evolutionary diversifications? Nematodes can help address such questions given their easy husbandry, simple diets, and rapid genetic and experimental manipulation, in particular in isogenic self-fertilizing hermaphrodites.

The developmentally plastic soil nematode *Allodiplogaster sudhausi* has several unique features, including its extreme body size (Fig. 1A, B) and predatory activity as juveniles and adults (*4*). As a distant relative of the model organism *Pristionchus pacificus* (*5*), it is one of the few self-fertilizing hermaphrodites outside *Pristionchus* with available CRISPR technology tools (*6*). *A. sudhausi* forms teeth-like denticles that were previously described to exhibit developmental plasticity with two discrete morphs, the narrow-mouthed ‘stenostomatous’ (St) and the wide-mouthed ‘eurystomatous’ (Eu) (*4*). In contrast to other predatory species, both mouth forms are able to kill other nematodes, including *C. elegans*. We recently showed *A. sudhausi* underwent a whole genome duplication (WGD) that is not seen in its closest relative *Allodiplogaster seani* (*6*). Additionally, *A. seani* is similar in size to *C. elegans* and *P. pacificus* (Fig. 1B), indicating that the extreme increase in body size of *A. sudhausi* correlates with the WGD event.

**Fig. 1.**
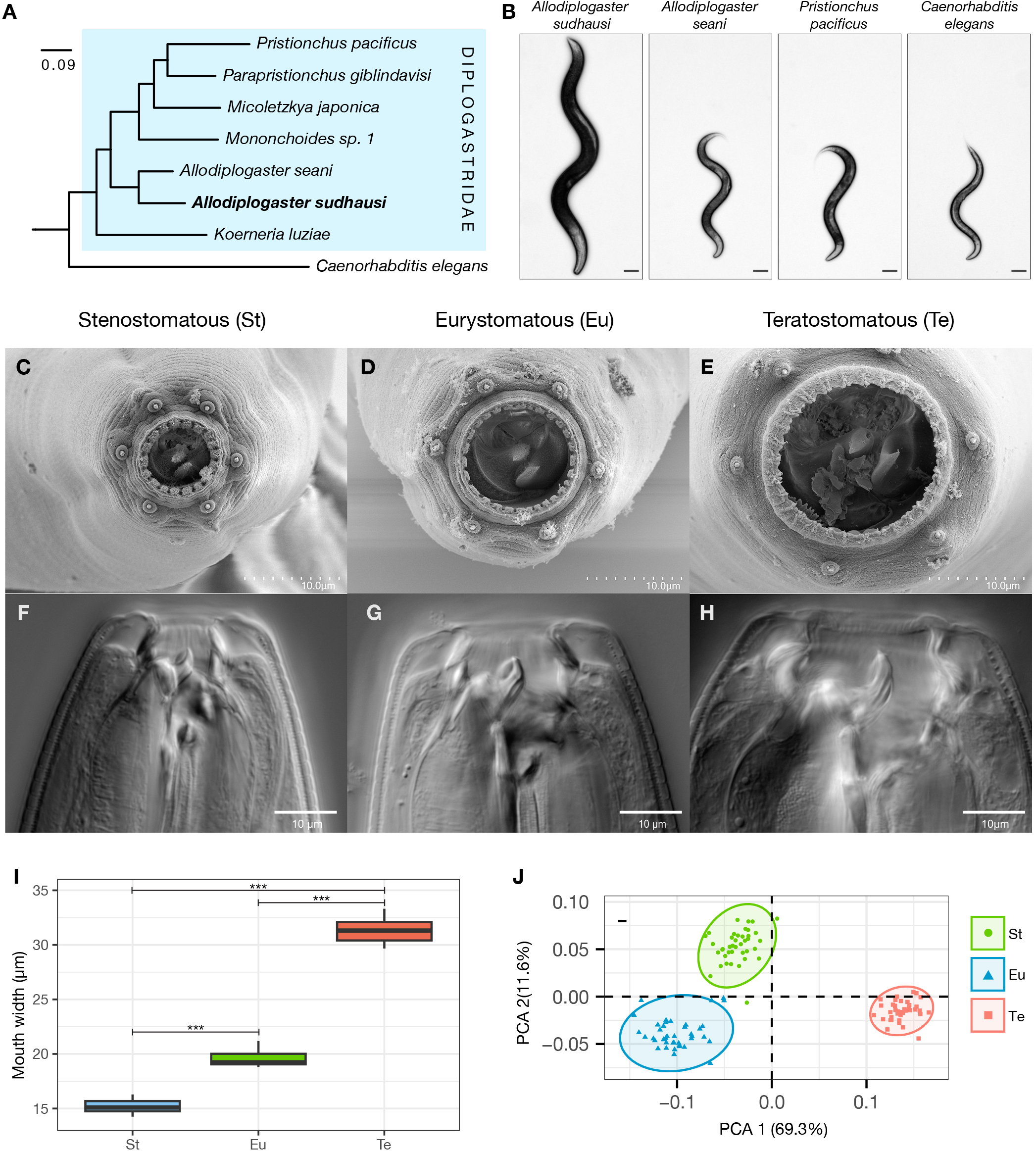
*A. sudhausi* forms an additional third mouth morph. **(A)** The phylogeny shows *Asudhausi* diverged early within the Diplogastridae. **(B)** The size of different hermaphrodite/female free-living nematodes is shown, with *A. sudhausi* much larger than others. **(C-E)** Scanning electron microscopy (SEM) pictures of *A. sudhausi* **(C)** St, **(D)**, Eu and **(E)** Te mouths, with the dorsal tooth on top. Differential interference contrast (DIC) microscopy pictures of *A. sudhausi* **(F)** St, **(G)**, Eu and **(H)** Te mouths, with the dorsal tooth on the left. **(I)** Mouth width measurements show significant size differences between St, Eu and Te mouths (Kruskal-Wallis, p < 0.001; Pairwise Wilcox test, p < 0.01 for each comparison) n = 8 - 12 biological replicates). **(J)** Geometric morphometrics of the three morphs show significant differences in their shape dimensions (pairwise PERMANOVA, p < 0.001 for each comparison) n = 40 for each morph.

### *A. sudhausi* exhibits three distinct mouth-forms

WGD is often associated with morphological novelties; however, it is unknown if it also affects developmentally plastic traits. To observe potential changes in mouth-form plasticity of *A. sudhausi*, we cultured these worms on a collection of bacterial, fungal and nematode diets. Remarkably, we found three distinct adult mouth-forms when *A. sudhausi* was grown on *Escherichia coli* OP50 bacteria, *C. elegans* N2 worms and *Penicillium camemberti* fungi (Fig. 1 C-H). Specifically, on *E. coli*, the standard maintenance method for nematodes, *A. sudhausi* formed the narrow St morph (Fig. 1C, F; fig. S1). On *C. elegans*, an inducer of Eu in other nematodes (*7*), *A. sudhausi* also became Eu (Fig. 1D, G). In contrast, *A. sudhausi* was able to form a third morph on *P. camemberti* (Fig. 1E, H). While the *E. coli* and *C. elegans* diets caused 100% St and Eu morph formation, respectively, the fungal diet resulted in both the St and third morph. This new morph was also seen at low frequency on starved *E. coli* plates (fig. S2). Measurements of the stoma (mouth width) and geometric morphometrics (*8*) of the mouth revealed significant differences between the three morphs in size and shape (Fig. 1I, J). As the third morph has the largest mouth, we termed it ‘teratostomatous’ (Te), from the Greek word *teras* for monster. Note that previous studies in *A. sudhausi* only observed two distinct morphs, but they did not test different dietary conditions (*4, 9*). Taken together, we have identified a third discrete mouth-form in *A. sudhausi*.

Next, we investigated whether the Te morph is restricted to *A. sudhausi* or if it is also found in other nematodes. We focused on the sister species *A. seani* and grew it under the same three dietary conditions. *A. seani* became St on *E. coli* and *P. camemberti*, and Eu on *C. elegans*; however, it did not form the Te morph (fig. S3). Additionally, we tested a range of different bacteria, fungi and other worms, but were unable to produce any Te equivalent on *A. seani*. Starved plates also did not contain the Te morph. Similarly, no other species of *Allodiplogaster* or the closely related genus *Koerneria* have been described to form three mouth morphs. Thus, the Te morph is only found in *A. sudhausi* and likely evolved *de novo* after the two *Allodiplogaster* species diverged. Together, these findings indicate that *A. sudhausi* shows three novel traits that are not present in its closest relative; WGD, increased body size and a third mouth-form.

### The Te morph is induced by poor nutrition, crowding and displays cannibalism

The induction of a third morph by *P. camemberti* was unexpected. Why would a fungus better known for the cheese it makes induce a new morph? We tested other fungi, including *Penicillium rubens, Penicillium digitatum, Emericella nidulans, Aspergillus* spp., and *Botrytis cinerea*, which all induced the Te morph (table S1). Therefore, Te morph induction is due to the fungal diet and not *P. camemberti* in particular. One potential reason might be the low nutritional value of these fungi, which could initiate a stress response in *A. sudhausi*. This would be consistent with our observation that the Te morph is seen on starved *E. coli* plates. Besides starvation, crowding also results in stressful environments in nematodes. Indeed, we found that crowding increased Te occurrences on *P. camemberti* (Fig. 2A). When few worms were present, the majority became St; however, higher density led to increased Te incidence. This trend was seen across differently sized plates, with smaller plates requiring less worms to reach the same Te percentage as larger plates (Fig. 2B). In contrast, crowding did not result in the formation of the Te morph on *E. coli* (fig. S4). These results suggest that stressful environmental conditions, combining crowding and low nutrition, induce the Te morph. Consistent with this, we show that Te animals have a smaller brood size and shorter body length than St and Eu animals (fig. S5). *A. sudhausi* therefore appears to have evolved a new phenotype in response to stressful conditions. However, what is the ecological significance of being Te?

**Fig. 2.**
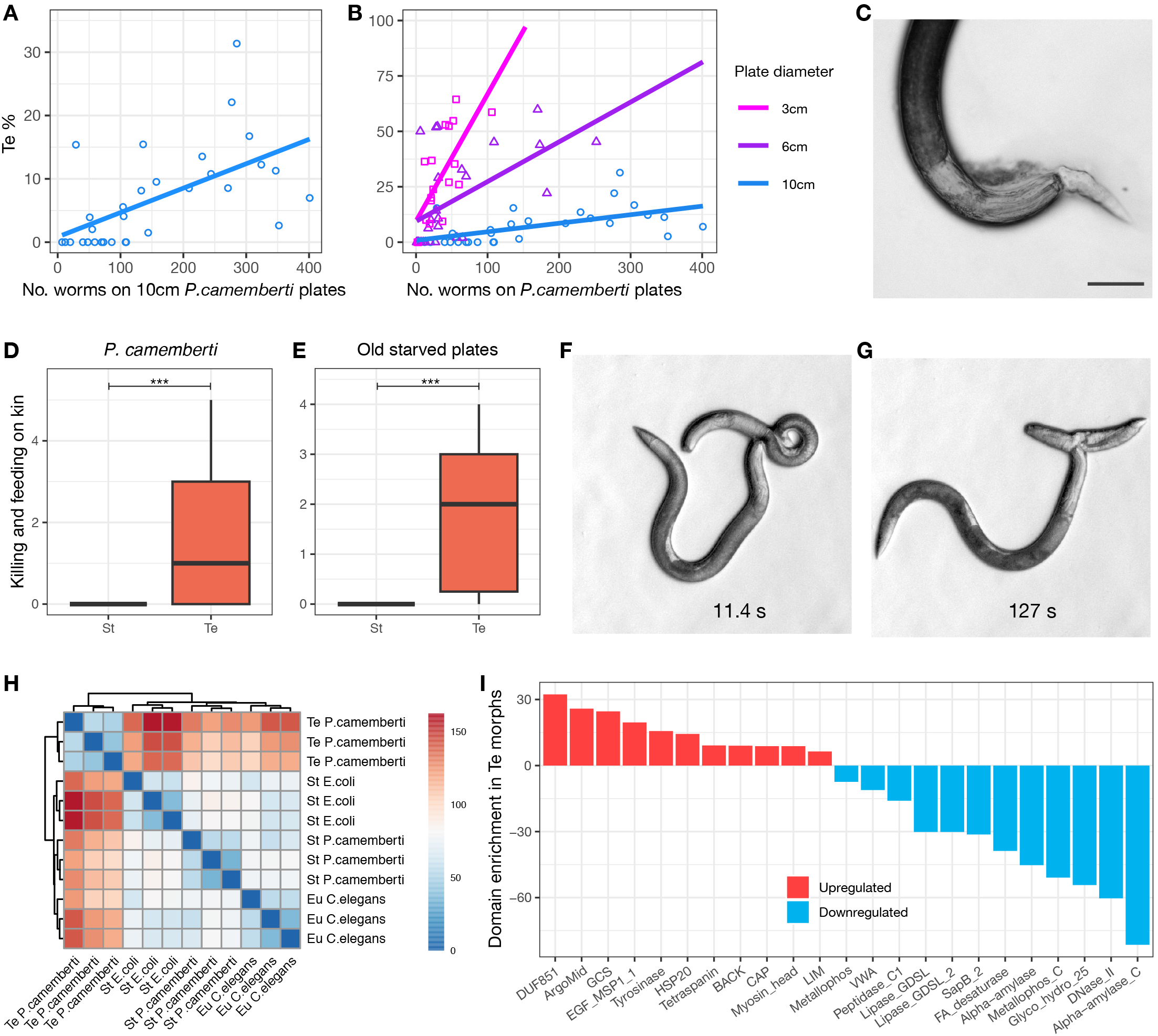
The Te morph cannibalizes, seemingly as a stress responses. **(A-B)** Crowding is positively correlated with Te morph induction. The number of adult *A. sudhausi* on **(A)** 10cm *P. camemberti* plates is associated with greater Te percentage (pearson’s product-moment correlation, p < 0.001) while **(B)** increased density increases Te induction as smaller plates show similar trends but at higher rates (pearson’s product-moment correlation, p < 0.01 for 6cm and p < 0.001 for 3cm). Linear regression lines are plotted for all. (C) An image shows Te feeding on younger kin (the tail of the juvenile is still seen), as commonly occurs. **(D)** From *P. camemberti* plates, Te morphs cannibalize on their living kin, unlike St which don’t kill and feed on kin at all (Mann-Whitney U test, p < 0.01). n = 20 each. **(E)** From old starved plates, only Te cannibalize (Mann-Whitney U test, p < 0.01). n = 14 each. **(F-G)** Te feed on other Te morphs and are quick at devouring prey, with pictures taken **(F)** 11.4 and **(G)** 127 seconds after feeding started. The full video can be found online (Movie S2). **(H) A** heat map of the adult transcriptome (measuring Euclidean distances between samples) shows that the Te morphs exhibit markedly distinct expression compared to the Eu and St samples, even from the same *P. camemberti* fungus. (I) Genes were identified that show significantly differential expression in Te versus other morphs (adj. p < 0.5, log2(FC) >2). The Pfam domains of these genes was predicted and the significantly enriched domains that are upregulated and downregulated in Te samples is depicted.

Stressful and low nutritional conditions induce cannibalism in a number of vertebrates and invertebrates (*2, 10*). In nematodes, previous studies in *P. pacificus* have shown morph-specific cannibalism (*11*). These studies also revealed that the likelihood of cannibalism negatively correlates with genetic relatedness of *P. pacificus* isolates and isogenic kin do not kill each other at all (*12*). In contrast, we observed regular cannibalism of the Te morph on their kin (Fig. 2C). Specifically, systematic studies using well-established predation assays (*13*) revealed that only the Te morph exhibits cannibalism against kin, whereas St and Eu animals did not (Fig. 2D,E; fig. S6). Although adult Te worms usually prey on juveniles (Movie S1), cannibalism against adults can also occur including on other adult Te animals. Cannibalism occurred in Te from both *P. camemberti* and starved plates (Fig. 2D,E). Movie S2 shows an adult Te that quickly digests another Te adult (Fig. 2F,G). Thus, cannibalism in *A. sudhausi* appears Te-specific and occurs despite the worms being isogenic. Therefore, our findings represent a novel type of cannibalism.

To determine if the new behaviour of Te adult morphs is associated with differential gene expression, we generated transcriptomes of adult worms of all three morphs. Specifically, we compared four distinct conditions; Eu animals from *C. elegans* cultures, St animals from *E. coli* and *P. camemberti* cultures and Te animals also from *P. camemberti* cultures. We found that Te morphs had markedly different expression patterns (Fig. 2H). In total, we identified 1841 differentially expressed genes that are shared in Te *vs*. the other three conditions (log2FC>2, adj. p<0.5) (Fig. 2I). Notably, genes containing a heat shock protein (HSP20) domain were significantly upregulated in Te morphs. As HSPs are commonly activated during stress response (*14, 15*), this finding further supports the idea that the Te morph is a response to low nutritional conditions resulting from starvation and crowding.

### Conserved sulfatase switch genes control morph-form regulation

Next, we wanted to know how mouth-form plasticity in *A. sudhausi* is genetically determined. In principle, the regulation of the Te morph may involve a novel regulatory network or the co-option of pre-existing machinery. To examine this, we looked at the *P. pacificus* mouth-form gene regulatory network, which has been well characterized (*16*) (Fig. 3A). A supergene locus plays a crucial role in *P. pacificus* mouth-form determination (*17, 18*). Specifically, the *eud-1/* sulfatase gene acts as a developmental switch with *eud-1* mutants constitutively St under all conditions (*19*). The *eud-1* switch gene is homologous to *C. elegans sul-2* and has two more paralogs in *P. pacificus*. However, only *Ppa-eud-1* is involved in mouth-form regulation, whereas mutants in the other two genes display wild type mouth forms. In *A. sudhausi*, four homologs of *Ppa-eud-1* have been identified, which are not clustered as in *P. pacificus* (*17, 20, 21*). These four genes represent two pairs resulting from the recent WGD. We refer to the one pair as *Asu-sul-2-A* and *Asu-sul-2-B*, and the other pair as *Asu-sul-2-D* and *Asu-sul-2-E* (Fig. 3B, table S2) (*20*).

**Fig. 3.**
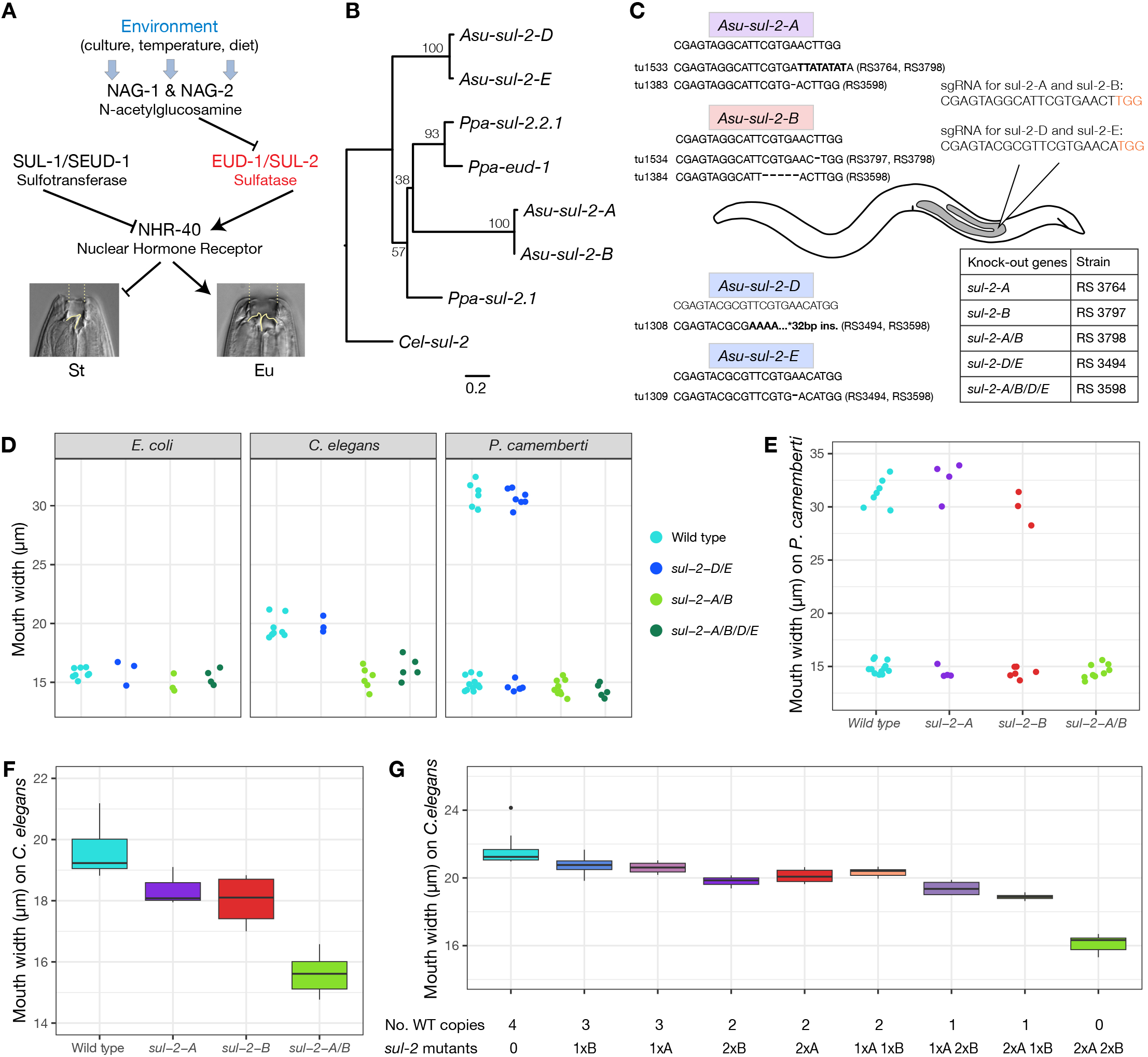
Genome editing indicates two functionally conserved sulfatases regulate both Eu and Te morphs. **(A)** Key genes that regulate mouth-form in *P. pacificus* are shown (17, 19, 22, 23). One of the major players, *eud-1*, encodes a sulfatase that regulates the Eu morph. **(B)** The phylogenetic relationship of the *eud-1* homologs in *P. pacificus* and *A. sudhausi* is shown, with C. *elegans sul-2* as the outgroup and bootstrap values indicating node support. There are four homologs in *A. sudhausi*, with two of each clustering, likely resulting from WGD. The relationship of the *A. sudhausi* sulfatase genes to *Ppa-eud-1* is not clear due to low bootstrap support. **(C)** The genotype of the *sul-2* mutant alleles is shown as well as the corresponding strains. One sgRNA was designed for each pair of genes. The PAM site next to the sgRNAs is indicated in red. **(D)** The mouth width of double and quadruple mutant lines was examined after being grown on the three conditions. Mutants with frameshift knock-outs in both *sul-2-A* and *sul-2-B* remain constitutively St under the three assays, in contrast to the *sul-2-D/E* mutant that displays wild type Eu and Te phenotypes on C. elegans and P. camemberti, respectively. n ≥ 14 biological replicates for each line. **(E)** The mouth width of knock-out mutants grown on *P. camemberti* is shown. The single mutants, *sul-2-A* and *sul-2-B*, are capable of becoming Te in contrast to the *sul-2-AIB* double and *sul-2-AIBIDIE* quadruple mutant lines. n ≥ 8 biological replicates for each line. **(F)** The mouth width of knock-out mutants grown on C. *elegans* is shown. Interestingly, the single mutants, *sul-2-A* and *sul-2-B* have an intermediate mouth width in comparison to wild type and *sul-2-A/B* mutants. **(G)** Crosses between wild type and mutants produced progeny with varying amount of wild type copies in *sul-2-A/B* when grown on C. *elegans*. There is a gradual decrease in mouth width as wild type copy number decreases and a stark drop in mouth width when there are no wild type copies. n ≥ 3 biological replicates for each line.

We targeted all four *A. sudhausi sul-2* homologs using CRISPR technology. Given the high sequence similarity of the WGD-derived gene pairs, we used identical sgRNAs for each and were able to generate single and double knockouts per pair (Fig. 3C). This strategy allowed us to obtain quadruple, double and single sulfatase gene knockouts. The quadruple mutant displayed the St morph under both *C. elegans* and *E. coli* diets, indicating that sulfatases regulate the Eu phenotype in *A. sudhausi*, identical to *P. pacificus* (Fig. 3D). Strikingly, when we grew the quadruple mutant on *P. camemberti*, all animals were St as well. Thus, both mouth-form decisions are regulated by *sul-2*/sulfatases.

### The Te morph is regulated by the same genes that control Eu formation

Next, we wanted to know if all sulfatases are involved in mouth-form regulation or if only one pair of sulfatase genes is functionally homologous to *Ppa-eud-1*. We found that the *Asu-sul-2-D/E* double mutant knockout displayed a wild type phenotype under all three dietary conditions, indicating this pair of genes has no role in mouth-form determination (Fig. 3D; fig S7). In contrast, the *Asu-sul-2-A/B* double knockout did not form either the Eu or Te morphs (Fig. 3D; table S3). Thus, the *Asusul-2-A/B* gene pair is functionally homologous to *Ppa-eud-*1 in term of Eu formation, and additionally controls the *A. sudhausi*-specific Te morph.

These findings culminate in the question as to whether *Asu-sul-2-A* and *Asu-sul-2-B* are redundant in mouth-form regulation. If both genes are fully redundant, single gene knockouts would be phenotypically wild type. In contrast, if *Asu-sul-2-A* and *Asu-sul-2-B* acquired different functions one would expect distinct phenotypes of single mutant animals. We found that single mutants displayed a wild type phenotype when grown on *P. camemberti* (Fig. 3E). This finding indicates that both genes are functionally redundant for Te morph formation.

In contrast, single mutants grown on *C. elegans* suggest more complex regulatory interactions involving dosage effects in Eu morph formation. Specifically, we found an intermediate mouth width of *Asu-sul-2-A* and *Asu-sul-2-B* single mutants when compared with wild type and double mutant animals (Fig. 3F). This surprising finding might indicate a quantitative role of *Asu-sul-2-A* and *Asu-sul-2-B* for the formation of the Eu phenotype. To test this hypothesis, we crossed wild type and mutant lines to generate progeny with four, three, two, one and zero wild type copies of *Asu-sul-2-A*/*B*. Phenotypic characterization of these individuals indicates a gradual decrease of mouth width from four wild type copies to one (Fig. 3G). Note the strong drop from one to zero wild type copies, suggesting that one functional copy of either gene is sufficient to recapitulate Eu-type features. These results support a dosage-dependent function of *Asu-sul-2-A*/*B* in the regulation of the Eu morph. Together, our findings indicate that *Asu-sul-2-A*/*B* control both Eu and Te formation with different regulatory mechanisms in both mouth-form decisions.

### Developmental plasticity and WGD facilitate morphological and behavioural diversity

In summary, the discovery of an additional mouth morph associated with cannibalistic behaviour supports the importance of developmental plasticity as a major driver of morphological and behavioural diversification. Our study identifies a pre-existing developmental switch gene that controls the Eu *vs*. St decision to then be co-opted for the regulation of the Te morph. The co-occurrence of three novel traits; WGD, doubling in body size and the evolution of a third mouth-form, are intriguing. We are currently unable to determine the exact order in which these characters have been gained, due to the absence of additional wild isolates and/or more closely related species. However, the WGD-derived *Asu-sul-2-A/B* already show functional divergence in Eu morph formation. Such molecular processes provide ecological opportunities highlighting the importance of plasticity. The cannibalistic Te morph, induced by crowding and starvation, enables greater chance of long-term survival in stressful conditions. This phenomenon relates to examples of cannibalism in salamanders, toads and fish (*2, 10*). However, *A. sudhausi* represents, to the best of our knowledge, the first animal system with regular cannibalism against genetically identical kin. Together, nematode mouth-form plasticity and predation provide a powerful system to investigate evolutionary innovation, its underlying molecular mechanisms, and behavioural and ecological consequences.

## Acknowledgments

We thank Eva Briem for capturing the movies. We thank Jürgen Berger for the SEM pictures. We also thank the Sommer Lab for advice and discussions, particularly Christian Rödelsperger.

## Author contributions

S.W. performed all experiments, with assistance from H.W. Data analysis was performed by S.W. Experiments were designed by S.W. and R.J.S. The manuscript was written by S.W. and R.J.S.

## Data and materials availability

Raw reads containing gene expression of adult morphs have been submitted to the European Nucleotide archive under the project accession number PRJEB65236. Remaining data is available in the supplementary materials.

## Materials and Methods

### Nematode maintenance and inbreeding

The *A. sudhausi* inbred strain SB413B/RS6132B (*6*) was used for wild type analyses. *A. seani* (RS1982) and *C. elegans* (N2) were also used. Mutant strains are frozen and kept at the Max Planck Institute for Biology. All nematode strains were maintained on nematode growth medium (NGM) agar plates with *E. coli* OP50 bacteria.

### Nematode culturing conditions

Freshly starved NGM *E. coli* plates containing many eggs were bleached (*24*) onto separate NGM containing three different food sources: 1) *E.coli*, 2) *C. elegans* and 3) *P. camemberti*. For *C. elegans*, plates with many larvae were washed with M9 and passed through a 20 µm nylon net filter (Merck) onto unseeded 6 cm NGM plates, so that only larvae passed through (*7*). *P. camemberti* was obtained from the Leipniz Institute DSMZ-German Collection of Microorganisms and Cell Cultures GmbH (https://www.dsmz.de) and maintained weekly on fresh unseeded NGM plates using sterile tips to transfer spores. Note that the other tested fungi were also obtained from DSMZ, except for *Aspergillus spp*. which was a lab contaminant that was naturally isolated. After bleaching of *A. sudhausi* onto these three conditions, all assays were maintained at 20°C until the eggs grew into adults. For all experiments, only young hermaphrodites were selected (containing less than five eggs).

### Count assays

The worms were counted and phenotyped under a Zeiss Stemi 508 light microscope when they reached adulthood. The mouth-form of adult worms was determined from all three assays. This data was then analyzed to compare the overall number of adult worms and the Te percentage.

### Standard Electron Microscopy (SEM)

Young adult worms were individually picked from *P. camemberti* and *C. elegans* culture conditions. St and Te adult hermaphrodites were taken from *P. camemberti* while Eu hermaphrodite were picked from *C. elegans*. The worms were moved into 1.5ml Eppendorf tubes containing 1ml PBS (Phosphate Buffered Saline) buffer. The SEM protocol was performed at the Electron Microscopy facility at the Max Planck Institute for Biology. The nematodes were fixed in 2.5% glutaraldehyde in PBS and post-fixed with 1% osmium tetroxide. Subsequently, samples were dehydrated in a graded ethanol series followed by critical point drying (Polaron) with CO2. Finally, the cells were sputter-coated with a 5 nm thick layer of platinum (CCU-010, Safematic) and examined with a field emission scanning electron microscope (Regulus 8230, Hitachi High Technologies).

### Microscopy images and measurements

Young adult hermaphrodites were fixed onto a solution with 5% Noble Agar and 0.3% NaN3, with M9 buffer for resuspension. They were imaged at 100x magnification using the Differential Interference Contrast (DIC) setting. Only worms fixed in the correct orientation (on the lateral side) were picked. Z-stacks were taken of the head region and stored as raw date czi files. Fiji/ImageJ (*25*) was used to measure the width of the mouth (stoma), using the base of the cheilostom and the start of the gymnostom the markers. All worms taken from the same assay plate were counted as one biological replicate.

### Worm size measurements

Young adult St and Te hermaphrodites from *P. camemberti* were moved to NGM plates without bacteria. Brightfield images were taken at 20x magnification using the Axio Zoom V16 microscope. Analysis was done on Fiji/ImageJ using the Wormsizer plugin for Image J/Fiji (*26*) to detect worm length. All worms from one assay plate were counted as one biological replicate.

### Geometric morphometrics

For each of the three morphs, young adult hermaphrodites were picked directly from their assay plates and image stacks were obtained using DIC microscopy, with the dorsal tooth always located on the left. Geometric morphometrics was performed using 18 fixed landmark coordinates, following the previous protocol (*27*). We generated landmark data for 40 individuals for each of the three morphs. Analysis was performed in RStudio (2021.09.2) (*28*) using the geomorph package (v. 4.0.1). Shape and form differences were visualized using PCA. To test for significant differences between morphs, we performed Procrustes ANOVA (PERMANOVA) using procD.lm. Pairwise PERMANOVA was then performed to determine differences between groups.

### Brood size analysis

Young virgin *A. sudhausi* hermaphrodites that had not yet laid eggs were moved from well-fed *E. coli* assay to NGM plates containing 30 µl litres of *E. coli*. They were transferred to fresh plates ever two days for a total of eight days. The number of live offspring that each hermaphrodite produced was counted. All plates were stored at 20°C. Progeny from adults belonging to the same plate were counted as one biological replicate.

### RNA sequencing and analysis

Young adult hermaphrodites were picked, moved into 20µl PBS buffer in 1.5ml Eppendorf tubes, and frozen at -80°C. RNA was extracted using Zymo Research Direct-zol RNA MiniPrep Kit following the manufacturer’s protocol, with the addition of flash freezing in liquid nitrogen three times after adding TRIzol, to ensure successful lysis. RNA was quality controlled using the NanoDrop One Spectrophotometer and sent to Novogene Co. for library prep and Illumina sequencing (paired end 150bp, 3G per sample). Over 20 million raw reads were generated for each sample. STAR (version 2.7.9a) (*29*) was used to align the reads against the genome (*6*), which were then summarized using featureCounts (v. 2.0.2) (*30*) (“-p” “-T” “8”). Subsequent analysis was performed using DesSeq2 (v. 1.38.3) (*31*). Low count data was filtered out (≥ 3 samples had a count of ≥ 10). For the heat map, data was log transformed (‘blind’ setting) and Euclidean distances were measured between samples. The R package pheatmap (v. 1.0.1) was used to generate a matrix showing sample distances. To examine the genes differentially expressed (DE) in Te morphs from *P. camemberti*, Te was compared to other morphs to determine DE genes that were upregulated (adj. p < 0.05, log2(FC) >2) and downregulated (adj. p < 0.05, log2(FC) < -2) in Te versus other conditions. The common DE genes upregulated in Te versus other morphs were then selected, as were the common DE genes downregulated in Te. Protein domains were annotated for all the genes using the Pfam database (v. 3.1b2) (*32*) as previously described (*6*). A Fisher’s 2x2 matrix was run, whereby each DE gene with a given domain was compared to the overall dataset to identify those that were significantly different (adj. p < 0.05, FDR). Enrichment values reflect occurrence of the significant DE domain-containing genes compared to overall genome-wide abundance.

### Feeding on kin assays

A feeding assay was adapted from that previously described (*7*): NGM was added to 12-well cell culture plates (Sarstedt). *A. sudhausi* juveniles were filtered into the wells using 41μm nylon net filters (Merck) so that only J3 and J2 larvae could pass through. Individual adult morphs were then added to each separate well containing younger kin. An Axio Zoom V16 microscope rotated between each well and took pictures every 20 seconds over a 90 minute period. Afterwards, the feeding events were counted. Videos S1 and S2 showing Te cannibalising was taken using an Axiocam attached to a Zeiss Stereo Discovery V20 microscope, set at 10 frames per second. The Axio zoom V16 microscope was used (Brightfield setting) to take the image of the adult Te feeding on a juvenile.

### Genetic crosses

*A. sudhausi* hermaphrodites at the late J4 stage were moved to separate plates containing *E. coli*. They were left for eight days on these plates, at which point they would have run out of sperm. They were then moved onto individual plates containing *E. coli* for 24 hours. Note that these plates were later checked to see if any viable progeny had in fact been produced. The old sperm-depleted hermaphrodites were then moved to separate plates containing 5µl of *E.coli* as well as 10 males of whichever line needed to be crossed. These worms were kept together for 24 hours to mate. They were then moved onto NGM plates containing *C. elegans* juveniles (as previously described). The resulting progeny would have one allele from the hermaphrodite and one from the male. Their adult mouth width was then measured. Note that these assays were performed for all crosses shown, even for hermaphrodites and males of the same line, so as to keep conditions consistent for comparisons.

### Phylogenetic trees

The species tree was previously used (*6, 9*). For the EUD-1/SUL-2 phylogeny, the protein sequence of *C.elegans* SUL-2 was obtained from wormbase.org (version WS284). The corresponding sequences for *P. pacificus* and *A. sudhausi* were found using prisitonchus.org. For *P. pacificus*, the annotation (El Paco V3 2020) for each protein is as follows, EUD-1: PPA43535, SUL2.2.1: PPA06135, SUL2.1: PPA21290. *A. sudhausi* gene models homologous to SUL-2 were also identified (Table S2). The phylogeny was generated using RAxML version 8.2.12 (raxmlHPC -f a -m PROTGAMMAAUTO -p 12345 -x 12345 -N 100), with maximum likelihood boostrap values included (raxmlHPC -f b -m PROTGAMMAILG) (13). The phylogeny was displayed using FigTree software (v.1.4.4) (tree.bio.ed.ac.uk/software/figtree).

### CRISPR Engineering and Mutant Identification

CRISPR knockouts were generated as previously described (*6*). CRISPR RNAs (crRNAs) and trans-activating crispr RNA (tracrRNA) (Cat. No. 1072534) were synthesized by Integrated DNA Technologies (IDT), while the Cas9 endonuclease (Cat. No. 1081058) was purchased from IDT. The CRISPR/Cas9 complex was prepared by mixing 0.5 mg/ml Cas9 nuclease, 0.1 mg/ml tracrRNA, and 0.056 mg/ml crRNA in the TE buffer followed by a 10-min incubation at 37° C. Microinjections were performed in late-stage J4 hermaphrodites using an Eppendorf microinjection system. The sgRNA sequence was designed just before an NGG PAM site and targeted the putative exon two of all *A. sudhausi sul-2* homologs (Fig. 3C). Specific primers were designed for all four of the genes. The primers for *sul-2-A* are: 5′-AGAATGTAGCCAGGCAAGC-3′ and 5′-CTCAGTCGACATGGAAAAGC -3′ (445 bp amplicon). For *sul-2-B*: 5′-ATTGCAGGATACGGCGACC -3′ and 5′-CTAATCTCGTCTATGCCGACG ′-3 (442 bp amplicon). For *sul-2-D*: 5’-ATGGTCGACGATCTTGGTAGCG -3’ and 5’-CCTGGAATGGAAGTCACACATGG - 3’ (745bp amplicon). For *sul-2-E*: 5’-GGTTTGATGACGCGTCATTGC -3’ and 5’-CGACCATGCCCGTGATATAGC -3’ (862bp amplicon). Polymerase chain reactions (PCRs) were run using these primers on the F1 of injected worms. Heterozygotes were identified via Sanger sequencing (GENEWIZ Germany GmbH) and homozygous mutants were obtained by self-fertilizing heterozygotes to generate homozygous knock-out mutants. One sgRNA was designed for each pair of similar genes. Frameshift mutants in *sul-2-A* (RS3764), *sul-2-B* (RS3797) and double frameshift mutants in *sul-2-A/B* (RS3798) were obtained after single injection of the *sul-2-A/B* sgRNA. The *sul-2-D/E* (RS3494) double mutant was obtained after a single injection as the gRNA knocked out both genes. In order to generate the quadruple mutant, the *sul-2-A/B* sgRNA was injected into the *sul-2-D/E* double mutant line. The resulting injection resulted in triple mutants in *sul-2-A/B/D* and *sul-2-A/B/E*. These triple mutants were crossed to each other to eventually obtain the *sul-2-A/B/D/E* quadruple mutant knockout line (RS3598).

### Statistical analysis

The mean of technical replicates was determined for each biological replicate, which was subsequently analysed. For mouth width and worm size measurements the number of technical replicates varied, as not all worms were in the correct orientation to get accurate measurements. To test for significant differences between groups, the Shapiro-Wilk test was first performed to determine whether groups displayed normal distribution. If so, a student’s t test was performed to compare the two groups. If the assumption of normality was violated, nonparametric tests were performed: Mann-Whitney U test for two comparisons and Kruskal-Wallis for group comparisons between groups, followed by the Pairwise Wilcox test. To analyse the association between independent and dependent variables, Pearson’s correlation coefficient was used. All statistics and analysis was performed using RStudio Statistical Software (2021.09.2).

### Supplementary Materials

Figs. S1 to S7

Tables S1 to S3

References (*24*–*32*)

Movies S1 to S2

Data S1 to S5

