## Supplementary Material for "Conserved switch genes regulate a novel cannibalistic morph after whole genome duplication"

**The PDF file includes:**

Figs. S1 to S7
Tables S1 to S3

**Other Supplementary Materials include the following:**

Movie S1 to S2
Data S1 to S5

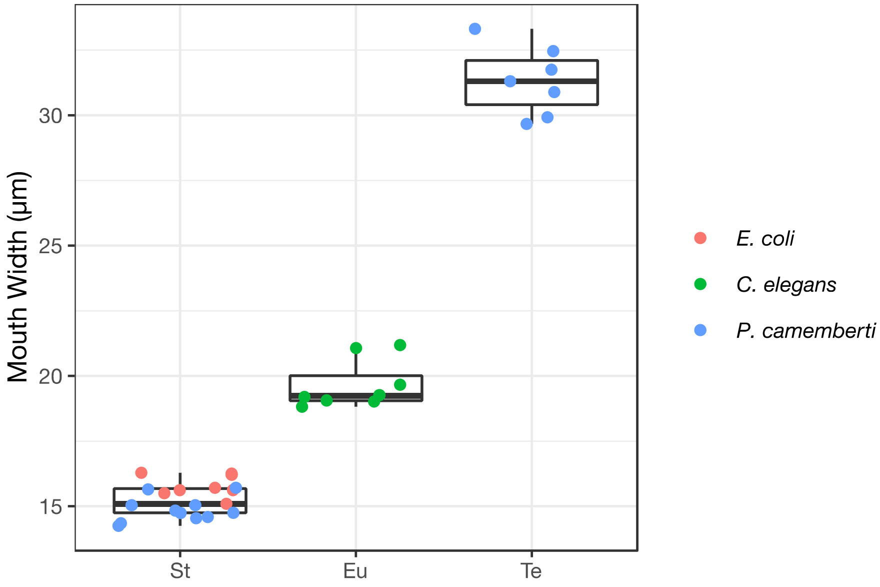


**Fig. S1.** The mouth width measurements of all three different assays is shown, with *E.coli* and *P.camemberti* constituting the St morph, *C.elegans* diets constituting the Eu and *P. camemberti* constituting the Te morph.

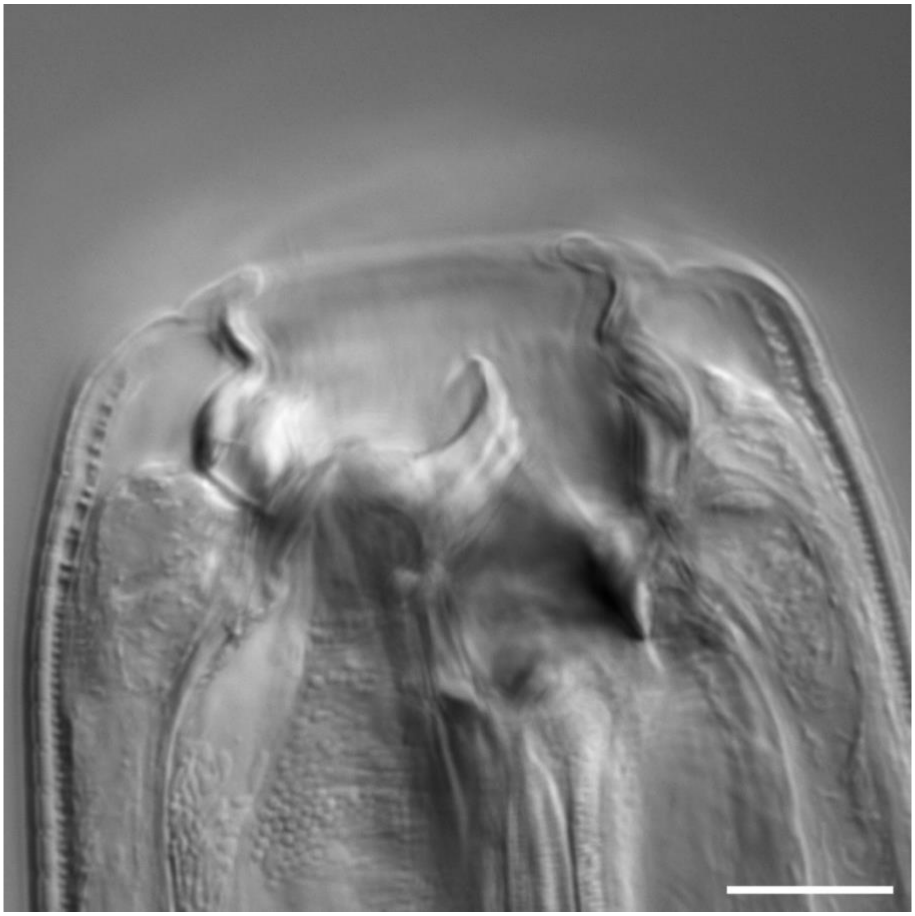

**Fig. S2.**

The Te morph can also be seen on old plates that have been starved of *E.coli*. Scale bar: 10 μm.

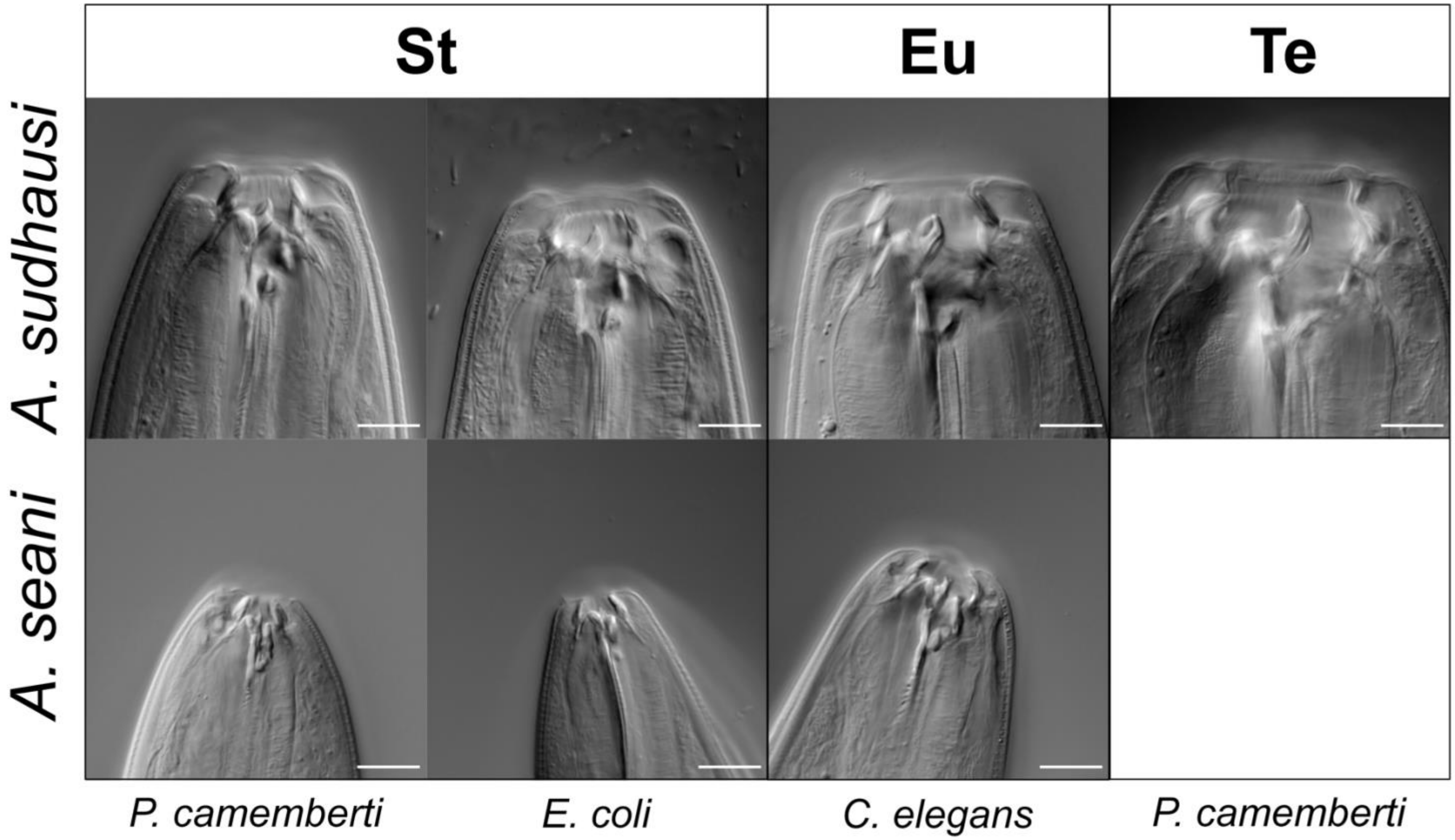


**Fig. S3.** *A. seani* can form the St (from *E.coli* and *P.camemberti*) and Eu (*from C.elegans*) morphs. No Te morph is seen from *P. camemberti* conditions. Scale bar: 10 μm.

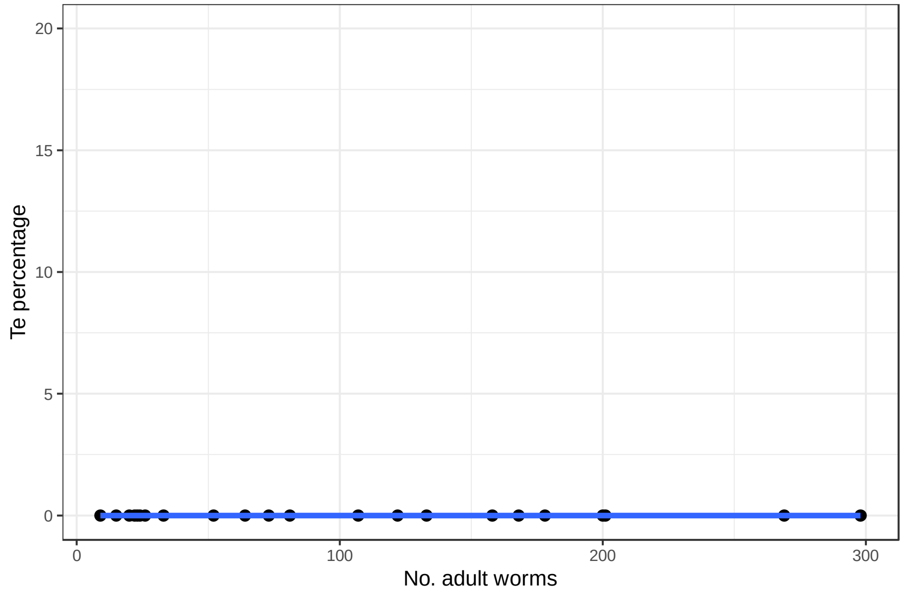


**Fig. S4.**

There is no correlation between the number of *A.sudhausi* adults and Te percentage. On *E.coli* plates, no worms form the Te morph. A linear regression plot is shown.

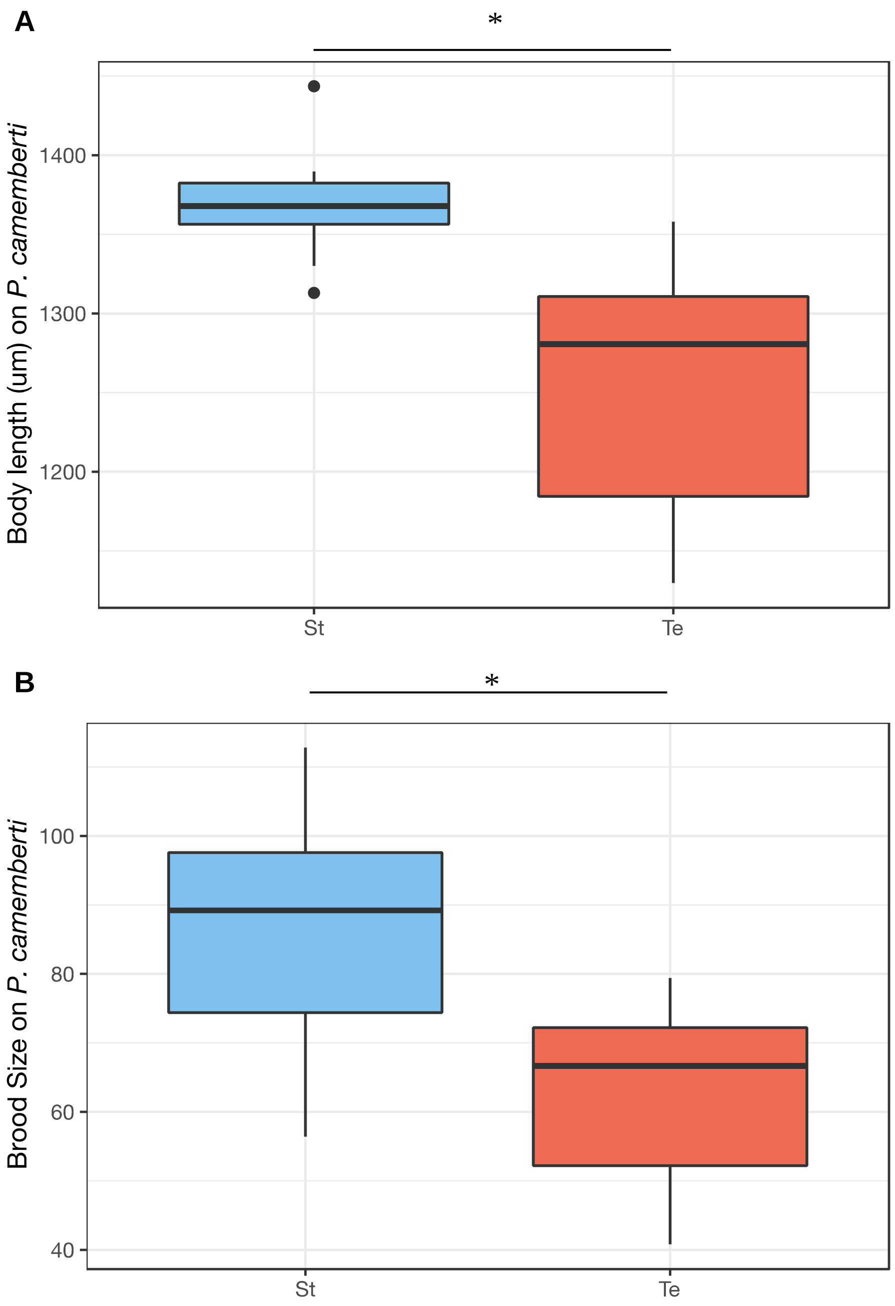


**Fig. S5.** Te morphs from *P. camemberti* assay show fitness costs compared to St morphs. (**A**) Te worms are significantly shorter than St (t test, $p < 0.05$). n = 10 biological replicates each. (**B**)Te worms also display decreased brood size (t test, $p < 0.05$). n = 7 biological replicates each.

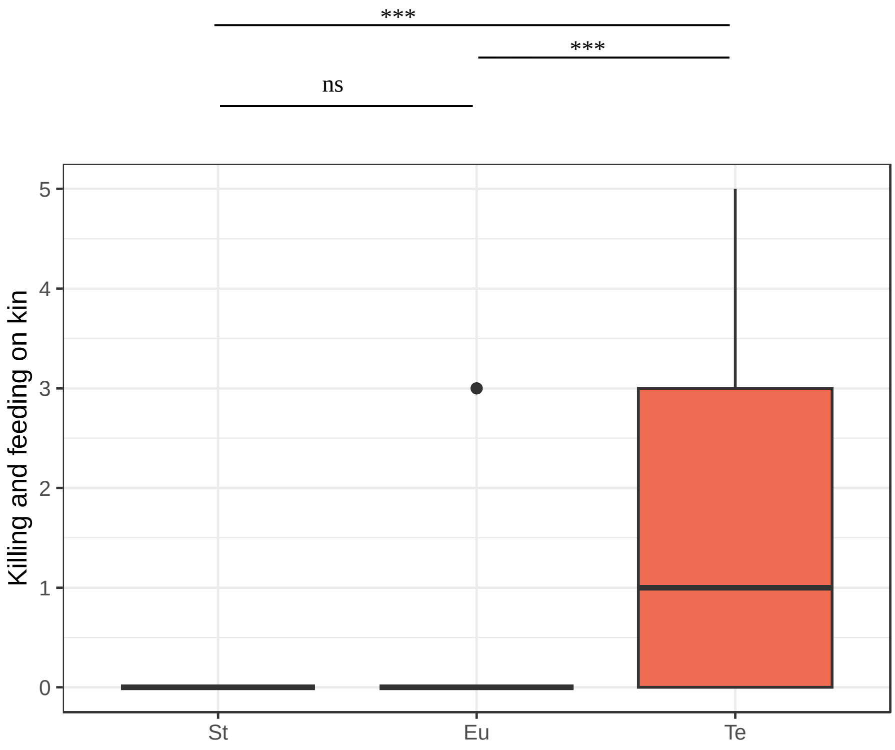


**Fig. S6.** Feeding of *A.sudhausi* on its younger kin was compared for all three mouth-form morphs. This includes St grown on *E. coli, P. camembert*i and old starved plates; Eu grown on *C. elegans,* and Te from *P. camemberti* and old starved plates. There were significant differences between Te and the other morphs (Kruskal-Wallis test, $p < 0.001$; Pairwise Wilcox, $p < 0.001$) but not between St and Eu ($p > 0.05$). The full dataset is shown in Data S4.

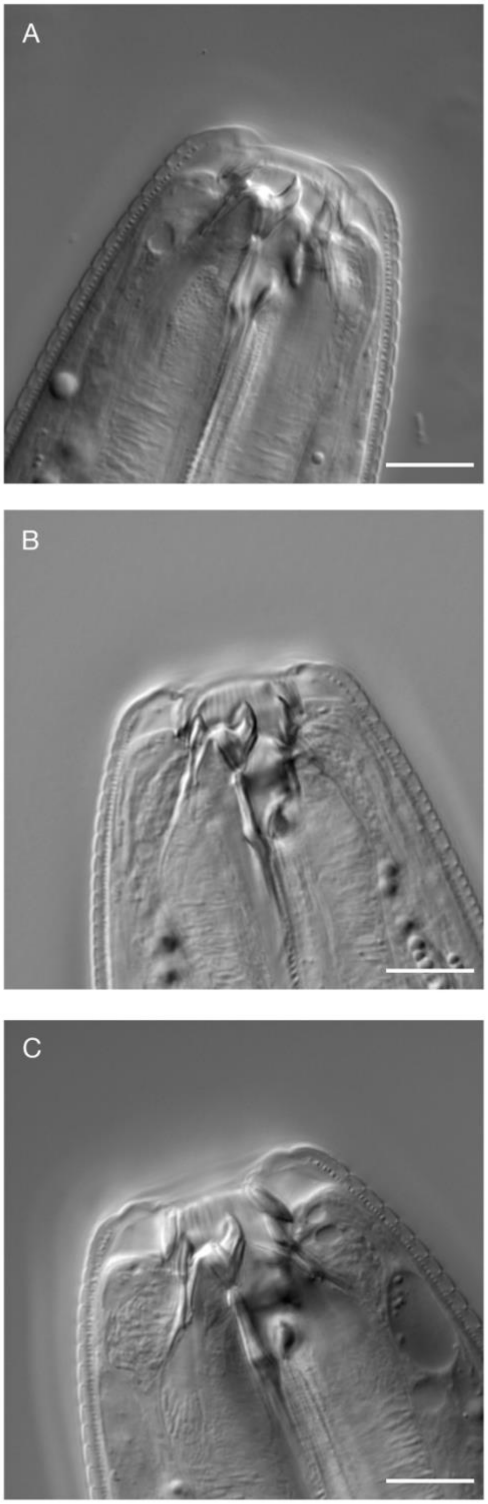


**Fig. S7.** DIC images of adult mouth-form in *sul-2-A/B* double mutants show they remain St when grown on (A) *E.coli*, (B) *C. elegans* and (C) *P.camemberti.* Scale bar: 10 μm.

**Table S1.** Table showing the different fungal strains *A. sudhausi* was initially tested on. All strains were able to produce the Te morph at varying percentages.

| **Fungus** | **Replicate** | **Te Percentage (%)** | **No. Hermaphrodites** |
|---|---|---|---|
| *Penicillium rubens* | 1 | 4,69 | 64 |
| *Emericella nidulans* | 1 | 2,27 | 88 |
| *Botrytis cinerea* | 1 | 52,38 | 21 |
| *Aspergillus spp.* | 1 | 54,55 | 22 |
| *Aspergillus spp.* | 2 | 57,14 | 28 |
| *Penicillium digitatum* | 1 | 7,14 | 42 |
| *Penicillium digitatum* | 2 | 5,26 | 38 |
| *Penicillium digitatum* | 3 | 17,39 | 23 |
| *Penicillium digitatum* | 4 | 12,90 | 31 |
| *Penicillium camemberti* | 1 | 20,31 | 64 |
| *Penicillium camemberti* | 2 | 8,14 | 86 |
| *Penicillium camemberti* | 3 | 4,08 | 49 |
| *Penicillium camemberti* | 4 | 9,52 | 84 |
| *Penicillium camemberti* | 5 | 10,68 | 103 |
| *Penicillium camemberti* | 6 | 8,51 | 94 |

**Table S2.** Table showing the gene names and their corresponding gene models. The matching gene and contig names in previous literature is also shown. Note that, based on our analysis, the previously annotated *sul-2-C* and *sul-2-D* are in fact one gene, which we refer to as *Asu-sul-2-D.*

| **Gene Name** | **Predicted Gene Model (Wighard et al. 2022)** | **Biddle & Ragsdale (2020)** | **Sieriebriennikov et al. (2018)** |
|---|---|---|---|
| *Asu-sul-2-A* | ALDISUDHAUS000034192 | *sul-2-A* | Contig 2291 |
| *Asu-sul-2-B* | ALDISUDHAUS000032677 | *sul-2-B* | Contig 5979 |
| *Asu-sul-2-D* | ALDISUDHAUS000002417 | *sul-2-C & sul-2-D* | Contig 474 |
| *Asu-sul-2-E* | ALDISUDHAUS000002604 | *sul-2-E* | Contig 8694 |

**Table S3.** Table showing the mouth-form of *sul-2A/B* and *sul-2-A/B/DE* adult mutants grown on *P. camemberti.* No Te morph was ever formed.

| Gene knock-out | Strain | Replicate | No. St | No. Te |
|---|---|---|---|---|
| *sul-2-A/B* | RS3798 | 1 | 56 | 0 |
| *sul-2-A/B* | RS3798 | 2 | 65 | 0 |
| *sul-2-A/B* | RS3798 | 3 | 6 | 0 |
| *sul-2-A/B* | RS3798 | 4 | 101 | 0 |
| *sul-2-A/B* | RS3798 | 5 | 133 | 0 |
| *sul-2-A/B* | RS3798 | 6 | 6 | 0 |
| *sul-2-A/B* | RS3798 | 7 | 73 | 0 |
| *sul-2-A/B* | RS3798 | 8 | 6 | 0 |
| *sul-2-A/B* | RS3798 | 9 | 165 | 0 |
| *sul-2-A/B* | RS3798 | 10 | 82 | 0 |
| *sul-2-A/B* | RS3798 | 11 | 27 | 0 |
| *sul-2-A/B/D/E* | RS3598 | 1 | 35 | 0 |
| *sul-2-A/B/D/E* | RS3598 | 2 | 8 | 0 |
| *sul-2-A/B/D/E* | RS3598 | 3 | 53 | 0 |
| *sul-2-A/B/D/E* | RS3598 | 3 | 75 | 0 |
| *sul-2-A/B/D/E* | RS3598 | 4 | 36 | 0 |
| *sul-2-A/B/D/E* | RS3598 | 5 | 89 | 0 |
| *sul-2-A/B/D/E* | RS3598 | 6 | 157 | 0 |
| *sul-2-A/B/D/E* | RS3598 | 7 | 186 | 0 |

**Movie S1. Te morph cannibalizes on younger kin.**
An adult Te hermaphrodite kills and feeds upon juvenile kin. Video set at 2x speed.

**Movie S2. Adult Te cannibalizes on another Te morph.**
An adult Te hermaphrodite latches to, kills and feeds upon another adult Te hermaphrodite. Video set at 2x speed.

**Data S1. (mouth_width_measurements.xlsx)**
Excel file containing the mouth width measurements of the wild type and mutant lines that were grown on *E. coli, C. elegans* and *P. camemberti.*

**Data S2. (worm_sizing.xlsx)**
Excel table showing the body measurements of wild type worms grown on *P. camemberti* using the WormSizer plugin. Mesaurements of young adult St and Te hermaphrodites are shown. The file displays volume, length, middle width, mean width and surface area (all in µm) of the worm body. The length measurements were compared in fig. S5A.

**Data S3. (brood_size_pcamemberti.xlsx)**
Excel table showing the number of viable progeny produced by wild type St and Te morphs that had grown up on *P. camemberti.*

**Data S4. (cannibalism.xlsx)**
Excel table displaying the number of juvenile kin fed on by different adult *A. sudhausi* morphs.

**Data S5. (GM_landmarks.txt)**
Text file containing the 18 landmark co-ordinates for the geometric morphometrics comparison between wild type morphs.
